# Presynaptic GABA_A_ Receptors Mediate Ethanol-Induced Suppression at a Central Auditory Synapse

**DOI:** 10.64898/2026.05.07.723441

**Authors:** Jana Nerlich, Sasa Jovanovic, Christian Keine, Stefan Hallermann, Ivan Milenkovic

**Affiliations:** Carl-Ludwig-Institute for Physiology, Faculty of Medicine, Leipzig University, 04103 Leipzig, Germany; Institute of Biology, Faculty of Life Sciences, Leipzig University; School of Medicine and Health Sciences, Department of Human Medicine, Carl von Ossietzky university of Oldenburg, 26129 Oldenburg, Germany; Research Center Neurosensory Science, Carl von Ossietzky Universität Oldenburg, Germany

## Abstract

Acute alcohol consumption impairs speech perception, particularly in noisy environments, but the synaptic mechanisms underlying this auditory deficit remain unknown. Here, we investigated how low-dose ethanol modulates the endbulb of Held synapse in the Mongolian gerbil cochlear nucleus, the first central relay of the auditory pathway. *In vivo* recordings showed that ethanol suppresses sound-evoked firing, an effect blocked by GABA_A_ receptor antagonists. Direct presynaptic patch-clamp recordings revealed that ethanol potentiates GABA_A_ currents at the endbulb of Held terminal but not at the postsynaptic spherical bushy cell. A high intra-terminal chloride concentration drives a GABA-mediated depolarization that activates low-threshold Kv1 potassium channels and shunts the presynaptic action potential. The reduced action potential amplitude limits calcium influx and glutamate release, effectively suppressing synaptic transmission. These findings identify ethanol-enhanced presynaptic shunting inhibition at the first central auditory synapse as a cellular mechanism for alcohol-induced auditory dysfunction.

## Introduction

Ethanol is one of the most widely consumed psychoactive substances and impairs cognition, motor control, and sensory perception (Oscar-Berman and Marinković, 2007; Field et al., 2010; Abrahao et al., 2017; Sharma et al., 2017; Rapp et al., 2022). Even at modest doses reached after a single alcoholic beverage (∼0.3‰ wt/vol; ∼6.8 mM), ethanol compromises auditory performance, particularly in complex acoustic environments (Choi et al., 2021). These deficits exemplify “cocktail party deafness”, the difficulty of understanding speech against competing background sounds (Cherry, 1953), to which acute ethanol consumption is one important contributing factor (Upile et al., 2007). In humans, low-dose ethanol reduces sensitivity to low-frequency speech cues and impairs speech-in-noise performance (Fitzpatrick and Eviatar, 1980; Pearson et al., 1999; Choi et al., 2021), compromising both social communication and the detection of auditory warning signals. Critically, these deficits originate from impaired central synaptic processing rather than peripheral signal transduction. Auditory brainstem responses (ABRs) recorded after acute ethanol ingestion show increased latencies in waves III-VII, which index successive brainstem processing stages, whereas waves I and II, reflecting auditory nerve activity, remain unchanged (Chu et al., 1978). Yet, the cellular mechanism by which low-dose ethanol disrupts central auditory function remains unidentified.

Behaviorally relevant ethanol concentrations (3–30 mM) can enhance tonic inhibition mediated by extrasynaptic GABA_A_ receptors (GABA_A_R) (Sundstrom-Poromaa et al., 2002; Wei et al., 2004; Olsen et al., 2007; Lobo and Harris, 2008). These receptors are persistently activated by ambient GABA in the nano-to-micromolar range (Farrant and Nusser, 2005; Brickley and Mody, 2012) and are particularly sensitive to low ethanol concentrations (Wallner et al., 2003; Santhakumar et al., 2007), making them prime candidates to mediate ethanol’s suppressive effect on central auditory function.

The endbulb of Held–spherical bushy cell (SBC) synapse in the cochlear nucleus is the first central relay of the auditory pathway, where large auditory nerve terminals faithfully transmit temporal information to downstream sound localization nuclei (Brawer and Morest, 1975; Sento and Ryugo, 1989; Blackburn and Sachs, 1990; Joris et al., 1994; Keine et al., 2017; Keine and Englitz, 2025). Inhibition at this synapse, although predominantly glycinergic (Wenthold et al., 1987; Wickesberg and Oertel, 1990; Nerlich et al., 2014b; Keine and Rübsamen, 2015), includes a functionally important GABAergic component (Nerlich et al., 2014b; Nerlich et al., 2017). Slow transmitter clearance and asynchronous release render this inhibition effectively tonic at physiological firing rates (Nerlich et al., 2014a; Nerlich et al., 2014b), providing an ambient GABAergic substrate on which low-dose ethanol could plausibly act and identifying the endbulb of Held – SBC synapse as a strong candidate for ethanol modulation.

To uncover the cellular mechanism by which low-dose ethanol disrupts central auditory processing, we investigated the Mongolian gerbil, a well-established model for human-like low-frequency hearing (Gleich and Struts, 2012; Heeringa and Köppl, 2022; Jüchter et al., 2022; Peterson et al., 2024), combining *in vivo* and *in vitro* recordings. Our results show that low-dose ethanol enhances presynaptic GABA_A_R-mediated shunting inhibition at the endbulb of Held terminal. This inhibition attenuates presynaptic action potential (AP) amplitude and consequently reduces glutamate release. These findings identify a cellular mechanism by which low-dose ethanol intake perturbs early auditory processing.

## Results

### Ethanol attenuates sound-evoked activity via GABA_A_ receptors

To determine how ethanol (EtOH) modulates auditory signal encoding at the endbulb of Held – SBC synapse, we recorded spontaneous and sound-evoked activity from SBCs while locally applying ethanol via iontophoresis through a piggy-back electrode (Havey and Caspary, 1980; Dietz et al., 2012; Keine et al., 2016). We used 40 mM ethanol in the iontophoresis pipette to compensate for the diffusion gradient across the ∼15–25 µm distance between the application and recording electrodes, aiming to effective tissue concentrations approximating blood and brain levels after a single alcoholic beverage (∼0.3‰ wt/vol; ∼6.8 mM) (Olsen et al., 2007).

Ethanol application did not alter spontaneous firing rates (Fig. 1B2, C1), indicating that basal synaptic transmission and spike generation were unaffected in the absence of sound (“Spont”, control = 22.5 ± 7.3 AP/s, EtOH = 22.1 ± 7.5 AP/s, p = 0.83, n=11, paired t-test). In contrast, ethanol significantly suppressed sound-evoked firing (Fig. 1B1–1B3), both at the characteristic frequency (CF) and across the entire frequency–response area (FRA) (Fig. 1C1; “CF”: control = 130.3 ± 13.6 AP/s, EtOH = 90.9 ± 11.1 AP/s, p = 0.01; “FRA”: control = 91.6 ± 11.6 AP/s; EtOH = 68.8 ± 10.3 AP/s, p = 0.009, n=11, paired t-test), while leaving the inhibitory sideband one octave above CF unchanged (control = 14.2 ± 5.4 AP/s, EtOH = 16.4 ± 7.1 AP/s, p = 0.55, n=11, two-way RM ANOVA).

**Fig. 1:**
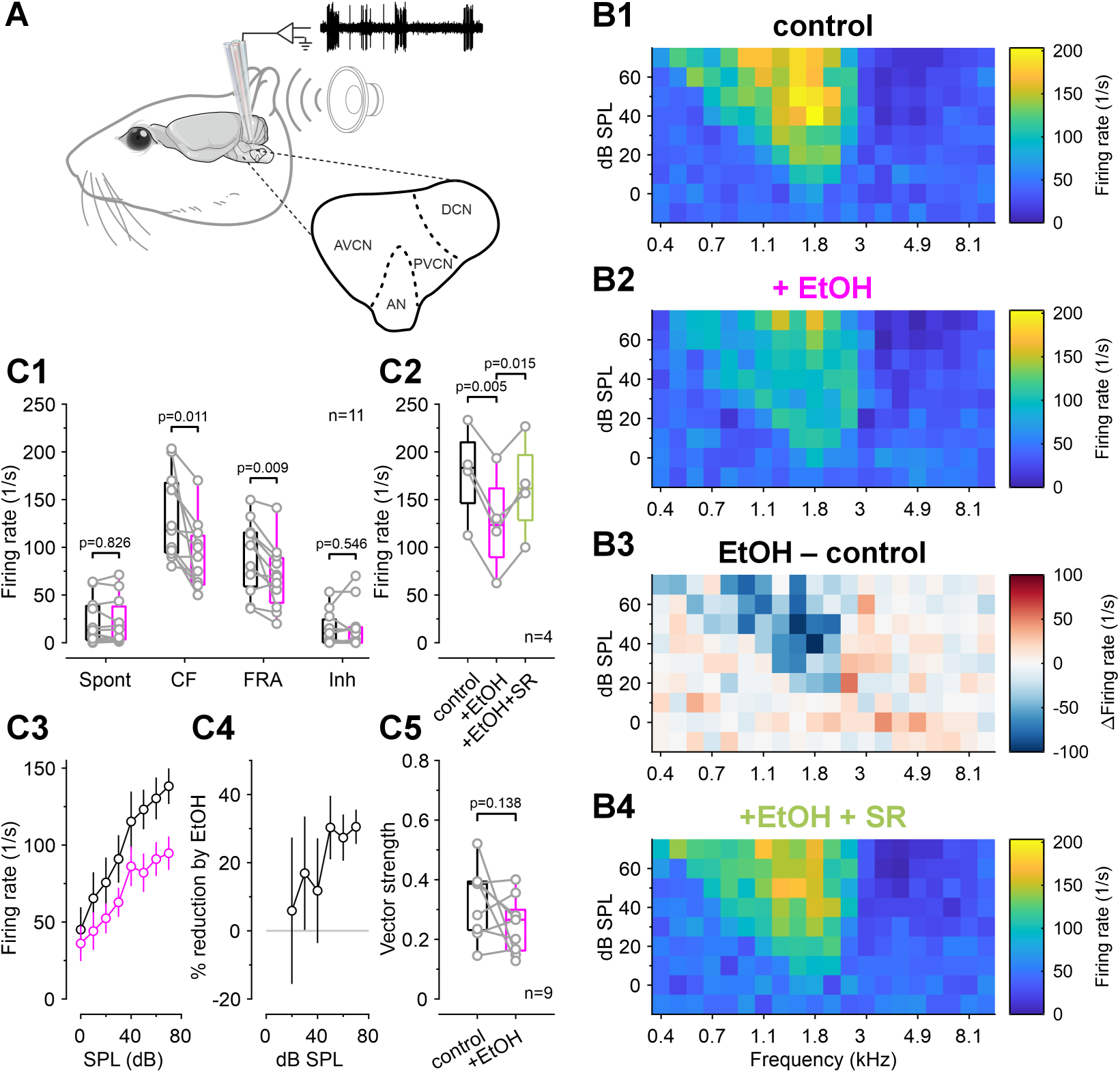
Ethanol suppresses sound-evoked activity in SBCs via GABA_A_R. (A) Schematic of the *in vivo* experimental setup. Multibarrel glass electrodes were used for juxtacellular recordings and iontophoresis in the rostral anteroventral cochlear nucleus (AVCN). DCN: dorsal cochlear nucleus; PVCN: posteroventral cochlear nucleus; AN: auditory nerve. (B) Representative frequency response areas (FRA) displaying firing rates (color scale) as a function of frequency and sound intensity. (B1) Control condition. (B2) Ethanol (EtOH) application reduces firing rates across the FRA. (B3) Difference map (EtOH – control) confirms broad suppression (blue). (B4) Co-application of the GABA_A_R antagonist SR95531 prevents ethanol-induced suppression. (C1) Quantification of firing rates. Ethanol (magenta) significantly reduced sound-evoked activity at the characteristic frequency (CF) and across the whole FRA, without altering spontaneous activity or in the inhibitory sideband (Inh). Box plots show median and interquartile range (IQR); whiskers extend to 1.5 x IQR. (C2) Blockade of GABA_A_R with SR95531 reverses the ethanol-induced reduction in firing rates. (C3-C4) Rate-level functions at CF (C3) and the corresponding percentage reduction (C4) show that ethanol suppresses firing across a broad range of sound levels, with robust effects at >50 dB SPL (C5) Temporal precision, quantified as vector strength, is unaffected by ethanol.

To test whether this suppression was mediated by GABA_A_R, a prime target of ethanol (Davies, 2003; Lobo and Harris, 2008), we co-applied the specific GABA_A_R antagonist SR95531. Blocking GABA_A_R abolished the ethanol-induced reduction and restored firing rates to control levels (Fig. 1B4, 1C2, EtOH vs SR: p = 0.015; SR vs ctrl: p = 0.16, n=4, RM ANOVA). To resolve how the firing rate depression at CF depends on sound intensity, we analyzed rate-level functions. Ethanol suppressed firing across a wide range of sound pressure levels (SPLs), with the strongest effects at intensities >50 dB SPL (Fig. 1C3, C4). This intensity-dependent compression suggests degraded sound intensity coding at SPLs relevant for speech perception.

Finally, we assessed temporal precision, a key factor for sound localization and speech processing (Zeng et al., 2005; Abrams et al., 2006). Despite robust rate suppression, ethanol did not alter temporal precision as quantified by vector strength (Goldberg and Brown, 1969) (Fig. 1C5, control vs. ethanol p = 0.138, n=9). Because postsynaptic inhibition typically enhances temporal precision in SBCs (Kuenzel et al., 2011; Keine and Rübsamen, 2015; Keine et al., 2016), the absence of such improvement under ethanol points to a mechanism distinct from canonical postsynaptic inhibitory modulation, which we next examined directly.

### Ethanol selectively potentiates presynaptic GABA_A_ currents

To identify the site of ethanol’s action, we first examined postsynaptic GABA_A_R using whole-cell recordings from SBCs in acute slices. Although pressure application of GABA evoked robust inward currents, bath application of ethanol (6.8 mM) did not change their amplitude (Fig. 2A-D; G_GABA_ = 38.9 ± 7.8 nS, G_GABA+EtOH_ = 41.3 ± 4.1 nS, p = 0.68, n=8, paired t-test). Thus, postsynaptic GABA_A_R are insensitive to this ethanol concentration and are unlikely to explain the firing rate suppression observed *in vivo*.

**Fig. 2:**
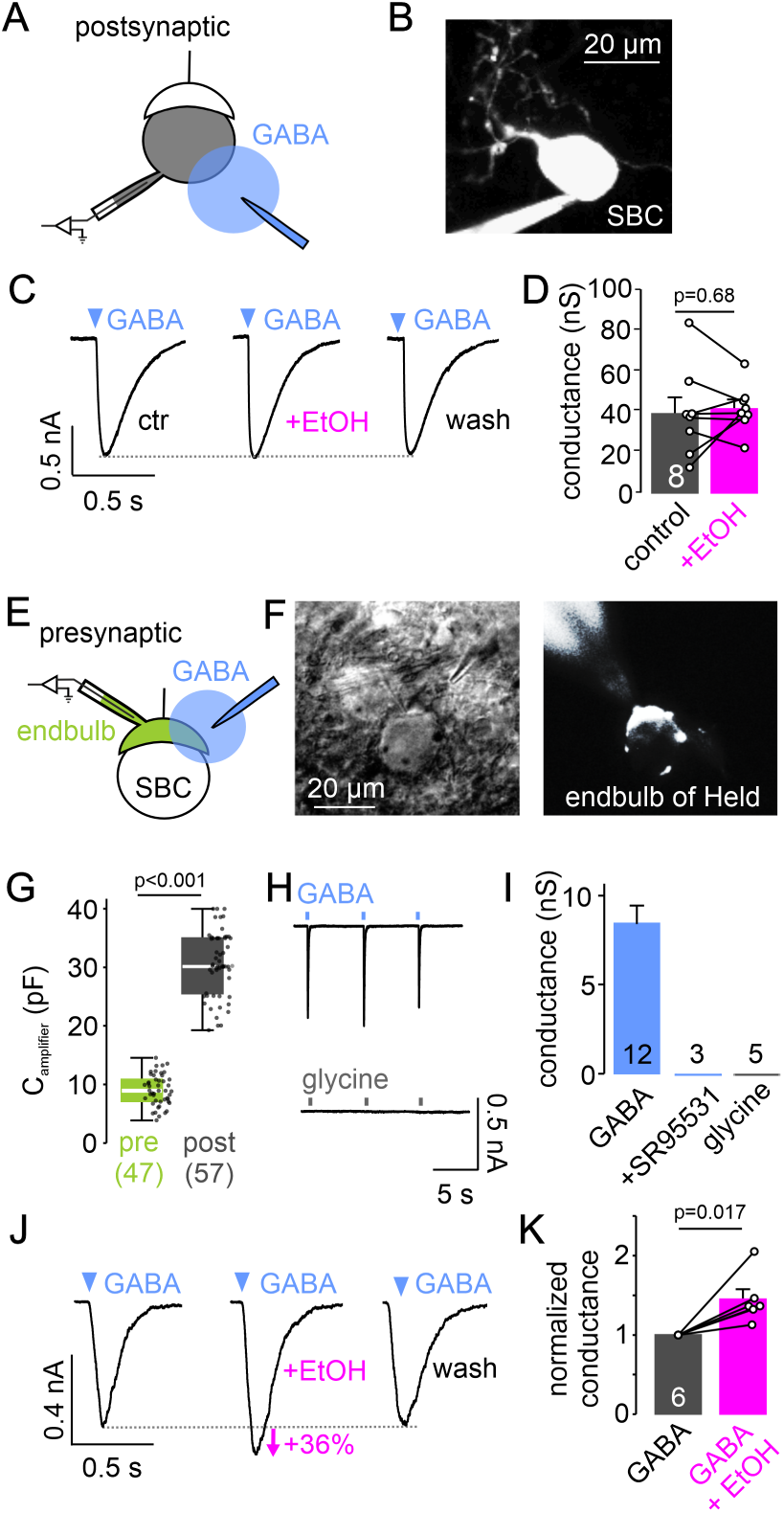
Ethanol selectively potentiates presynaptic GABA_A_R at the endbulb of Held. (A) Experimental configuration for whole-cell recordings from SBCs. GABA (500 µM) was pressure-applied near the soma. (B) Fluorescence image of a patch-clamped SBC filled with ATTO 488. (C) Representative GABA-evoked postsynaptic currents in control ACSF, during the bath application of ethanol (6.8 mM), and after washout. (D) Ethanol had no effect on postsynaptic GABA conductance (p = 0.68, n=8, paired t-test). Data points represent individual cells. (E) Configuration for presynaptic recordings from endbulb of Held terminals. (F) Differential interference contrast (left) and fluorescence (right) images of a recorded endbulb terminal enwrapping an SBC soma. (G) Membrane capacitance (C_amplifier_) clearly distinguishes presynaptic endbulbs (green) from postsynaptic SBCs (gray). Box plots show median and IQR, whiskers extend to 1.5 x IQR. (H) Presynaptic currents were evoked by local application of GABA but not glycine (both 500 µM). (I) Summary of pharmacological characterization. The GABA_A_R antagonist SR95531 (20 µM) abolished GABA-evoked responses, confirming GABA_A_R activation, while glycine had no effect. (J) Representative traces showing reversible potentiation of presynaptic GABA current by ethanol (6.8 mM). (K) Summary data for ethanol-induced enhancement of GABAergic conductance (paired t-test, n=6). Data are normalized to control. Cell numbers are indicated on the graphs.

We therefore investigated presynaptic receptors by performing direct whole-cell recordings from the endbulb of Held terminals (Fig. 2E). Presynaptic identity was confirmed by multiple criteria. Morphologically, terminals exhibited the characteristic endbulb structure, visualized live with the fluorescent dye ATTO 488 in the intracellular solution (Fig. 2F). Electrophysiologically, terminals displayed significantly smaller membrane capacitance than postsynaptic SBCs (Fig. 2G; C_amplifier_ endbulb = 9.0 ± 0.4 pF, n=47, C_amplifier_ SBC = 30.5 ± 0.7 pF, p < 0.001, n=57, t-test), higher input resistance (Lin et al., 2011), and a more hyperpolarized resting membrane potential (Fig. S1; R_input_ endbulb = 221.1 ± 10.6 MΩ, n=32, R_input_ SBC = 75.1 ± 11.3 MΩ, n=10, V_rest_ endbulb = −75.5 ± 1.1 mV, n=16, V_rest_ SBC = −66.7 ± 0.6 mV, p < 0.001, n=11, t-test).

Local application of GABA (500 µM, 10 ms, n=12) elicited inward currents in the presynaptic terminal (Fig. 2H, I; G_GABA_ = 8.4 ± 1 nS, n=12), whereas glycine (500 µM, 10 ms, n=5) had no effect. These GABA-evoked currents were abolished by SR95531 (n=3) (Fig. 2I), confirming that they are mediated by presynaptic GABA_A_R. In contrast to the postsynaptic site, bath application of ethanol (6.8 mM) significantly potentiated presynaptic GABA currents by 45 ± 13 % (Fig. 2J, K; p = 0.017, n=6, paired t-test). Together, these data demonstrate that ethanol selectively enhances GABA_A_R-mediated currents at the presynaptic terminal.

### Presynaptic GABA_A_ activation drives terminal depolarization

To determine the physiological impact of presynaptic GABA_A_R currents, we assessed the chloride driving force with gramicidin-perforated patch recordings from the endbulb of Held terminals (Fig. 3A), which preserve native intracellular chloride concentration (Ebihara et al., 1995; Akaike, 1996). Local GABA application at different holding potentials elicited currents that reversed at E_GABA_ = −31.1 ± 2.7 mV (n=11, Fig. 3B-D). This reversal potential was consistent across different buffering conditions (Fig. 3D; bicarbonate vs. HEPES p = 0.18, two-way ANOVA) and pipette solutions (E_GABA_ 7 mM vs. 150 mM [Cl⁻]_pip_ p = 0.3, two-way ANOVA), confirming the integrity of the perforated patch. Applying the Nernst equation (see Methods) indicates a native intraterminal chloride concentration of approximately 39 mM, considerably higher than in typical mature neurons, but consistent with high presynaptic chloride at the calyx of Held (Price and Trussell, 2006).

**Fig. 3:**
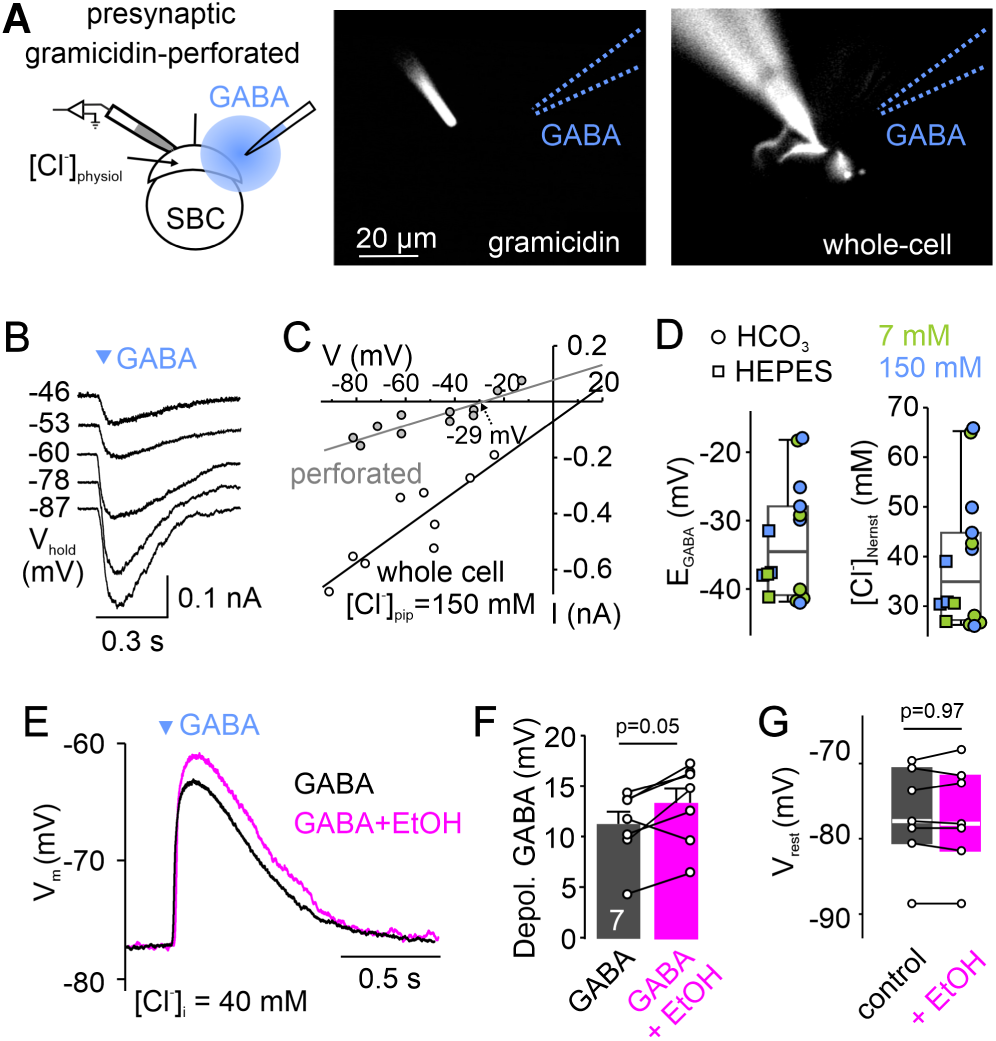
Presynaptic GABA_A_ receptors depolarize endbulb of Held terminals via chloride efflux. (A) Left: Experimental design for presynaptic gramicidin-perforated patch recordings with local pressure application of GABA. Middle/Right: Fluorescent images (ATTO 488) confirming patch integrity with the dye constrained to the pipette (middle) and subsequent transition to whole-cell configuration with the dye filling the terminal (right). (B) Representative GABA-evoked currents recorded at indicated holding potentials (V_hold_) in perforated patch configuration. (C) Current-voltage plots comparing GABA currents recorded in perforated patch configuration (gray symbols) versus whole-cell configuration with high internal chloride ([Cl^−^]_i_ = 150 mM, open symbols). (D) Population data for E_GABA_ (left) and calculated physiological intra-terminal chloride concentration ([Cl^−^]_Nernst_, right) (n=16). Symbols denote recording condition: HCO_3-_ (circle, n=11) versus HEPES-buffered ACSF (square, n=5), and 7 mM (green, n=7) versus 150 mM (blue, n=9) pipette chloride concentration. (E) Example current-clamp traces showing GABA-induced depolarization in control (black) and ethanol (6.8 mM, magenta) conditions. Recordings were obtained in whole-cell configuration with [Cl^−^]_pip_ = 40 mM, to match the native intraterminal [Cl^−^] determined in D (see Methods for details). (F-G) Population data: ethanol potentiates GABA-induced depolarizations (F; n=7, paired t-test), without altering resting membrane potential (G; n=7, paired t-test).

Due to the depolarized E_GABA_ relative to the resting membrane potential, GABA_A_R mediated a chloride efflux in perforated patch recordings, depolarizing the terminal by 14.0 ± 1.4 mV (n=13, Fig. S2). Comparable depolarizations were elicited in whole-cell configuration using 40 mM intra-terminal chloride (Fig. S2; [Cl⁻]_pip_ = 40 mM: V_rest_ = −74.4 ± 1.1 mV, ΔV_GABA_ = 10.7 ± 0.5 mV, n=20; perforated vs. whole-cell: V_rest_ p = 0.71, ΔV_GABA_ p = 0.25, two-way RM ANOVA), validating this configuration for subsequent experiments. Consistent with the voltage-clamp data, bath application of ethanol (6.8 mM) increased the GABA-induced depolarization by approximately 20% (Fig. 3E, F; p = 0.05, n=7, paired t-test) without altering the resting membrane potential (Fig. 3G; p = 0.97, n=7, paired t-test). Thus, presynaptic GABA_A_Rs mediate a depolarizing chloride efflux that is potentiated by low concentrations of ethanol, consistent with allosteric receptor modulation (Olsen, 2018).

### GABAergic depolarization attenuates presynaptic action potentials via Kv1 channels

The magnitude of presynaptic depolarization we observed lies within the range expected to recruit voltage-gated potassium channels at the endbulb, prompting us to ask whether GABA-induced depolarization affects the presynaptic AP. To address this question, presynaptic current-clamp recordings were performed, and APs were either evoked by stimulating the auditory nerve at 0.1 Hz, or by applying spike trains derived from *in vivo* recordings. Depolarization of the terminal, induced either by GABA application or by current injection mimicking the GABA-induced voltage shift (Fig. 4A-D), significantly reduced AP amplitudes (44%–50%) across all stimulation paradigms (Fig. 4C-E; effect of depolarization p < 0.001, effect of stimulation paradigm p = 0.009, two-way ANOVA), without altering AP duration (Fig. 4G). The extent of AP attenuation correlated strongly with the magnitude of depolarization (Fig. 4F; r = −0.85, p < 0.001, Spearman correlation) and occurred even with mild voltage changes induced by low GABA concentrations (Fig. 4F, S7D; 100 µM, ΔVm = 5.0 ± 0.3 mV, AP amplitude reduction: 12.7 ± 2.1 %, p = 0.002, n=6, paired t-test). These findings demonstrate that GABA-induced depolarization of the endbulb attenuates presynaptic AP amplitude in a dose-dependent manner.

**Fig. 4.**
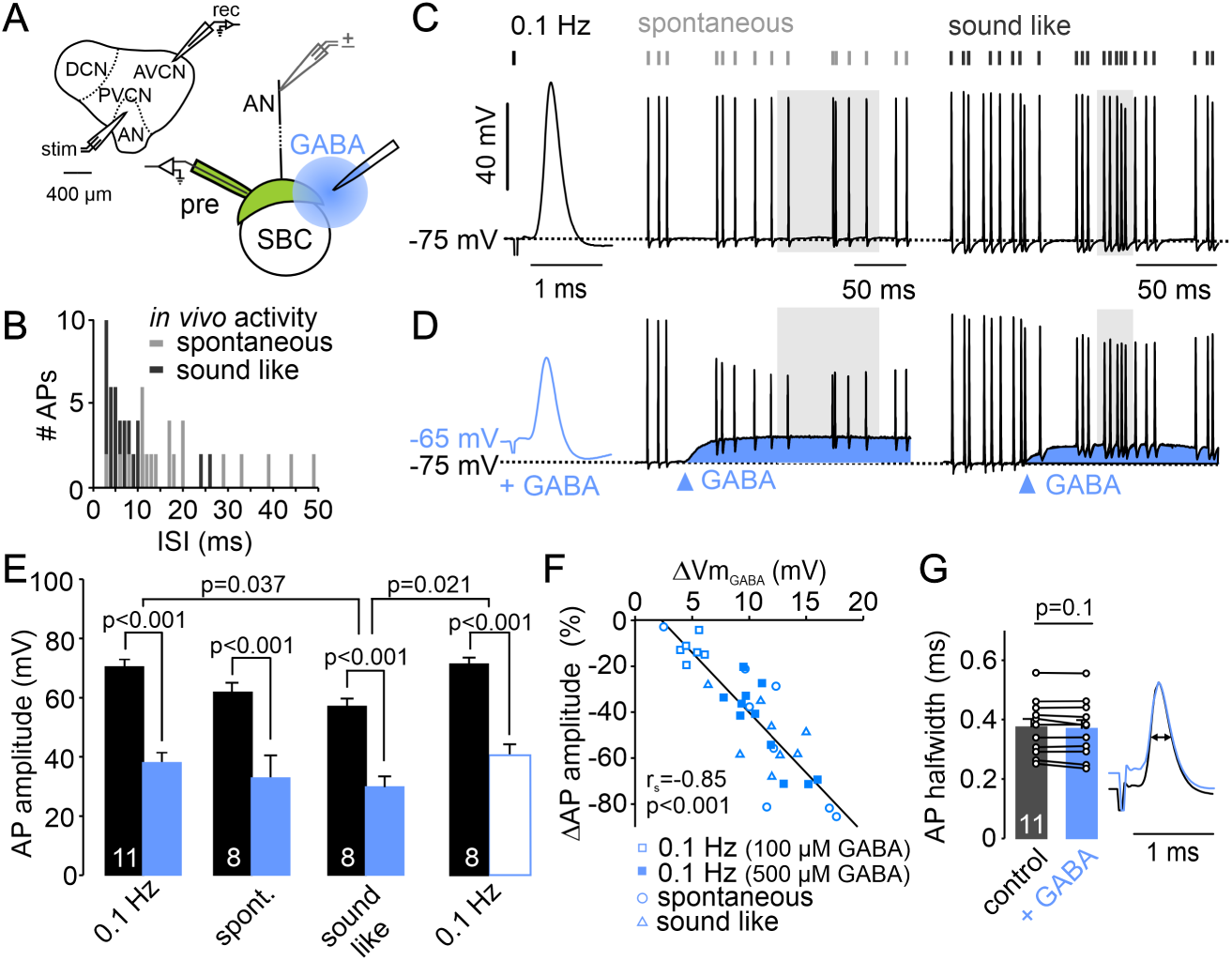
GABA-induced depolarization attenuates presynaptic action potentials. (A) Schematic of the experimental setup. Presynaptic APs were evoked via auditory nerve stimulation and recorded from the endbulb terminal in whole-cell configuration ([Cl^−^]_pip_ = 40 mM). GABA was applied locally via pressure application. (B) Inter-spike interval (ISI) histograms of the stimulus trains used to mimic spontaneous (gray) and sound-evoked (black) activity, derived from *in-vivo* SBC recordings. (C-D) Representative AP traces evoked by low-frequency (0.1 Hz), spontaneous, and sound-evoked stimulation in control conditions (C) and during GABA application (D). Blue shading indicates the membrane depolarization induced by GABA. Gray windows indicate the periods used for comparison. (E) Summary of AP amplitudes. GABA significantly reduces AP height across all stimulation paradigms (blue bars). Direct current injection (white bar, right) mimicking the GABA induced voltage shift, produced comparable AP attenuation. (F) AP attenuation correlates strongly with the magnitude of membrane depolarization (ΔV_m_). Different symbols represent stimulation paradigms (open = 100 µM GABA, filled = 500 µM GABA). (G) GABA did not alter AP half-width (n=11, paired t-test), as illustrated by peak scaled example APs.

Low-voltage-activated potassium channels (Kv1) shape the AP waveform and ensure temporal precision and are therefore critical for controlling the firing pattern and timing accuracy of neurons (Dodson et al., 2003; Rothman and Manis, 2003; Johnston et al., 2010; Brown and Kaczmarek, 2011; Hoppa et al., 2014). Because Kv1 channels activate near the resting membrane potential and exhibit only weak inactivation (Brew and Forsythe, 1995; Rothman and Manis, 2003; Cao et al., 2007), we hypothesized that GABA-induced depolarization would recruit them and thereby shunt the presynaptic AP. To test this, we evoked presynaptic APs by current injection, yielding amplitudes and durations comparable to those elicited by auditory nerve stimulation (Fig. S3). Pharmacological blockade of Kv1.1/1.2 channels with dendrotoxin-K (DTX-K; 50 nM) induced repetitive firing, significantly broadened APs, and increased their amplitude without altering the resting membrane potential (Fig. 5A-D; AP half-width p = 0.002, AP amplitude p = 0.002, V_rest_ p = 0.89, n=8, one-way ANOVA). Critically, DTX-K treatment significantly diminished the GABA-induced reduction in AP amplitude, despite comparable levels of depolarization (Fig. 5E, F; AP amplitude: DTX-K = 88.3 ± 3.5 mV, DTX-K+GABA = 68.4 ± 5.5 mV, p < 0.001, n=8, paired t-test; effect size comparison: p = 0.025; depolarization: GABA = 11.3 ± 1.5 mV, n=6, DTX-K+GABA =12.1 ± 1.1 mV, p = 0.69, n=8, t-test). In contrast, blockade of high-voltage-activated Kv3 channels with tetraethylammonium (TEA) prolonged AP duration without affecting AP amplitude, while increasing input resistance (Fig. 5A-D; AP half-width p = 0.007, AP amplitude p = 0.97, R_input_ p < 0.001, n=10, one-way ANOVA). Together, these results indicate that GABA-induced depolarization attenuates AP amplitude through activation of Kv1 channels at the endbulb.

**Fig. 5.**
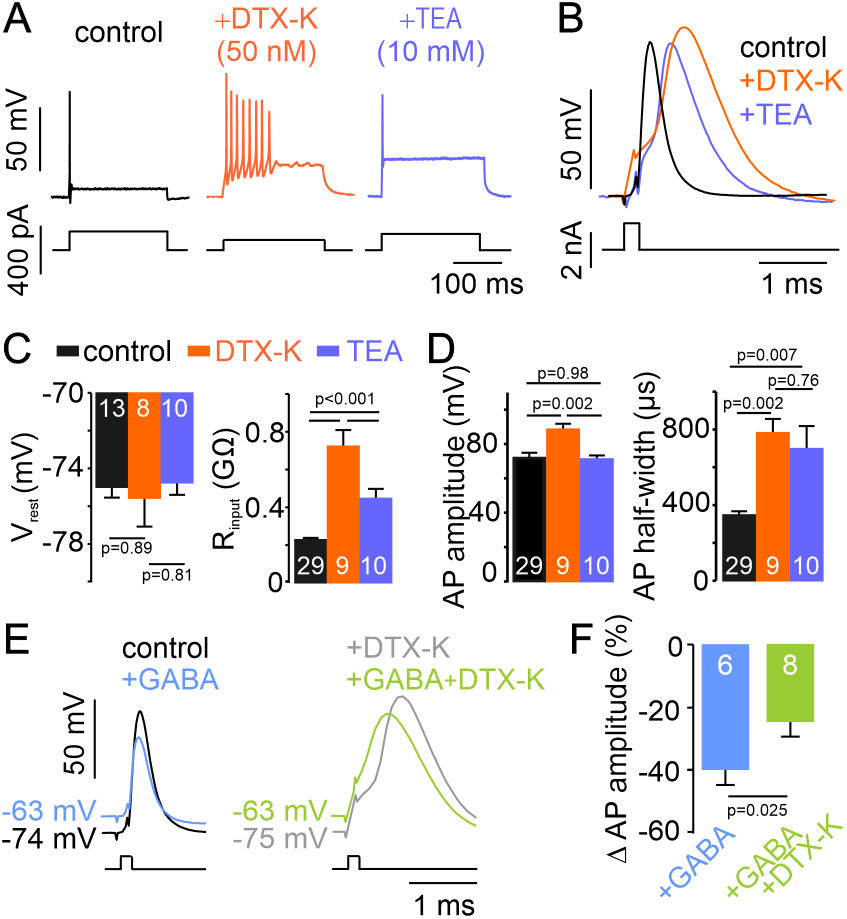
GABA-induced depolarization attenuates presynaptic APs via recruitment of Kv1 channels. (A) Representative voltage responses to 200 ms depolarizing current steps. Blockade of low-voltage-activated Kv1 channels with dendrotoxin (DTX-K, 50 nM, orange) induces repetitive firing. Blockade of high-voltage-activated Kv3 channels with TEA (10 mM, blue) preserves single spiking. (B) Overlay of single APs evoked by brief current injection (0.1 ms) showing waveform changes under channel blockade. (C) Resting membrane potential (V_rest_) was unaffected by K^+^-channels blockers (left), while input resistance (R_input_) was significantly increased, particularly by DTX-K (right). (D) Impact of channel blockade on AP shape. DTX-K selectively increased AP amplitude (left), whereas both DTX-K and TEA significantly prolonged AP half-width (right). (E) Representative APs recorded in control ACSF (left) or in the presence of DTX-K (right), before (black/gray) and during GABA application (blue/green). Note that the GABA-induced reduction in AP amplitude is markedly blunted when Kv1 channels are blocked. (F) Quantification of the GABA-induced reduction in AP amplitude (ΔAP amplitude). The suppressive effect of GABA is significantly attenuated in the presence of DTX-K, indicating that GABA-induced depolarization recruits Kv1 channels to shunt the spike. Cell numbers are indicated within the bars.

### AP attenuation reduces presynaptic calcium influx and glutamate release

Presynaptic AP amplitude determines calcium influx and neurotransmitter release at high-fidelity calyceal synapses (Borst and Sakmann, 1998; Lin et al., 2011; Young and Veeraraghavan, 2021). To determine whether the GABA-induced attenuation of the presynaptic AP compromises neurotransmitter release, we performed paired recordings from the endbulb terminal and the postsynaptic SBC (Fig. 6A). Previously recorded AP waveforms under control condition and during GABA application served as voltage-clamp commands; the GABA-like waveform reproduced both the reduced AP amplitude and the 10 mV baseline depolarization. The GABA-like waveform evoked markedly smaller presynaptic Ca²⁺ currents and total Ca²⁺ charge (Fig. 6B-E; I_Ca(v)_ control AP = 525.5 ± 43.7 pA, I_Ca(v)_ GABA-like AP = 268.9 ± 32.9 pA, Q_Ca(v)_ control AP = 193.6 ± 19.6 pC, Q_Ca(v)_ GABA-like AP = 106.0 ± 12.4 pC, p < 0.001, n=11, paired t-test), corresponding to a 46 ± 2.8 % reduction in presynaptic Ca^2+^-current (Fig. 6G; p < 0.001, n=7, paired t-test).

**Fig. 6.**
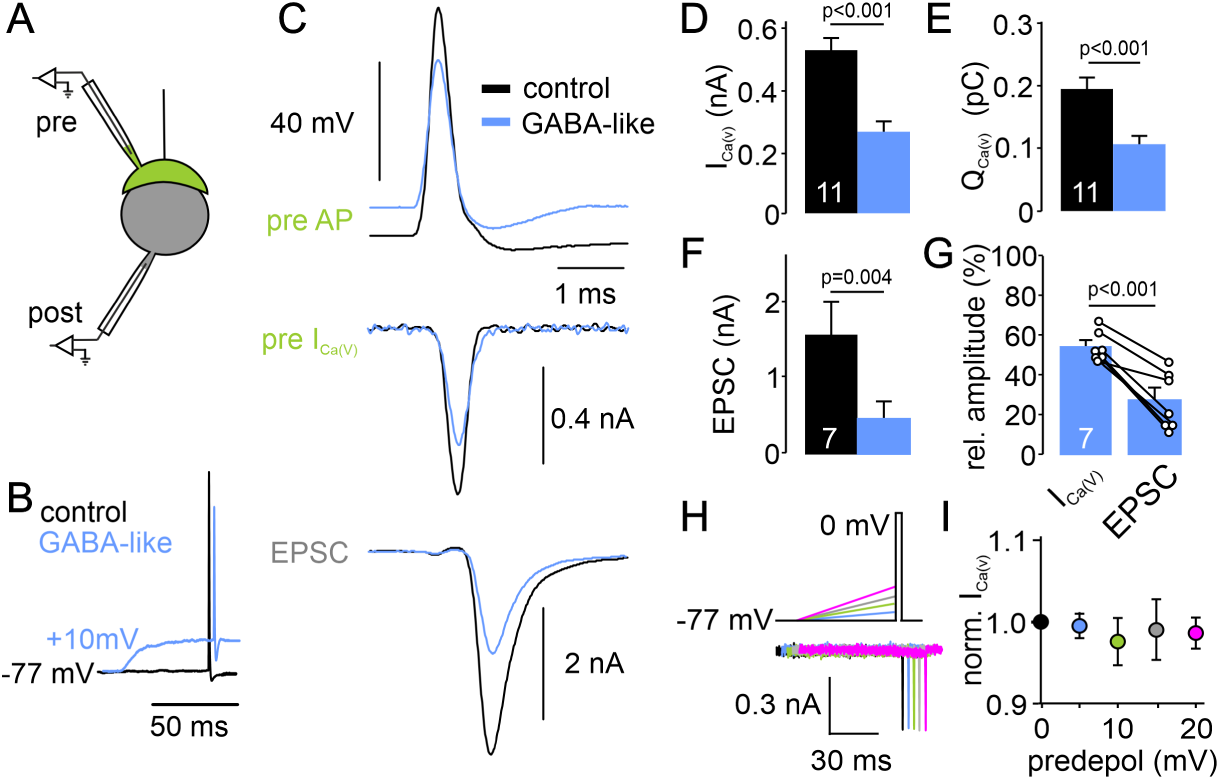
GABA-induced AP attenuation suppresses calcium influx and glutamate release. (A) Paired whole-cell recording configuration from a presynaptic endbulb and its postsynaptic SBC. (B) Voltage-command templates used to evoke presynaptic Ca^2+^-currents. The GABA-like waveform (blue) incorporated the AP attenuation and membrane depolarization observed during GABA application (see Fig. 4). (C) Representative trace from a paired recording. Top: Presynaptic voltage commands (control vs GABA-like). Middle: Evoked presynaptic Ca^2+^-currents (I_Ca(V)_). Bottom: Resulting excitatory postsynaptic currents. Both I_Ca(V)_ and EPSC amplitude are reduced by the GABA-like waveform. (D-E) Summary of presynaptic Ca^2+^-current amplitude (D) and charge transfer (Q_Ca(V)_) (E). The GABA-like waveform significantly reduced calcium entry (n=11, paired t-test). (F) Summary of EPSC amplitudes recorded from SBCs (n=7, paired t test). (G) Relative reduction in Ca^2+^-current versus EPSC amplitude. The suppression of release significantly exceeds the reduction in calcium influx, consistent with a supralinear relationship between Ca^2+^ entry and vesicle fusion. (H) Voltage protocol to test for Ca^2+^-channel inactivation. A square pulse to 0 mV (0.3 ms) was preceded by depolarization ramps of increasing amplitude (ΔVm = 0 – 20 mV, 70 ms, from V_hold_ = −77 mV). (I) Normalized I_Ca(V)_ amplitude plotted against pre-depolarization level. Ramp depolarizations did not alter Ca^2+^ currents(p = 0.7, n=6, RM-ANOVA).

Corresponding excitatory postsynaptic currents (EPSCs) were reduced even more strongly (Fig. 6C, F-G; EPSC control AP = 1.58 ± 0.44 nA, GABA-like AP = 0.46 ± 0.23 nA, 74 ± 5.3 % decrease, p = 0.004, n=7, paired t-test), consistent with the supralinear dependence of vesicular release at other high-fidelity synapses (Bollmann et al., 2000; Schneggenburger and Neher, 2000; Sakaba and Neher, 2001; Eshra et al., 2021). To test whether smaller presynaptic depolarizations than those evoked by 500 µM GABA also impair synaptic transmission, we repeated the paired-recording protocol using either a 5 mV current-injection depolarization or low-dose GABA (100 µM). Both manipulations significantly reduced EPSC amplitudes (Fig. S7B, C–F), indicating that mild presynaptic depolarizations, as expected from ambient GABA levels *in vivo*, compromise glutamate release. We next tested whether the reduced Ca^2+^ current reflects voltage-dependent inactivation of presynaptic Ca^2+^ channels due to the sustained 10 mV depolarization, rather than a consequence of reduced AP amplitude per se. Ca^2+^ currents were elicited with brief voltage steps to 0 mV preceded by depolarizing ramps of increasing amplitude. Pre-depolarization up to 20 mV did not change Ca^2+^ current amplitudes (Fig. 6H, I; I_Ca(v)_: 557–588 pA across ramps, p = 0.7; n=6; one-way RM ANOVA), arguing against Ca^2+^-channel inactivation as the underlying mechanism. Together, these results demonstrate that even modest GABA-induced depolarizations attenuate presynaptic AP amplitude and thereby reduce calcium entry and EPSC amplitudes.

### Ethanol potentiates presynaptic GABA_A_R-mediated synaptic suppression

Our findings demonstrate that ethanol enhances presynaptic GABA-induced depolarization (Fig. 3), which attenuates AP amplitude and curtails calcium influx (Fig. 4, 6). To directly assess the impact on glutamatergic transmission, we measured EPSCs evoked by electrical stimulation of the auditory nerve (50 pulses at 100 Hz) during transient GABA application via a puff-pipette (Fig. 7A). Presynaptic effects were isolated by voltage-clamping each SBC at its experimentally determined E_GABA_, thereby eliminating postsynaptic GABAergic currents (examples in Fig. 7B1, S4B, C at E_GABA_ = −63 mV).

**Fig. 7.**
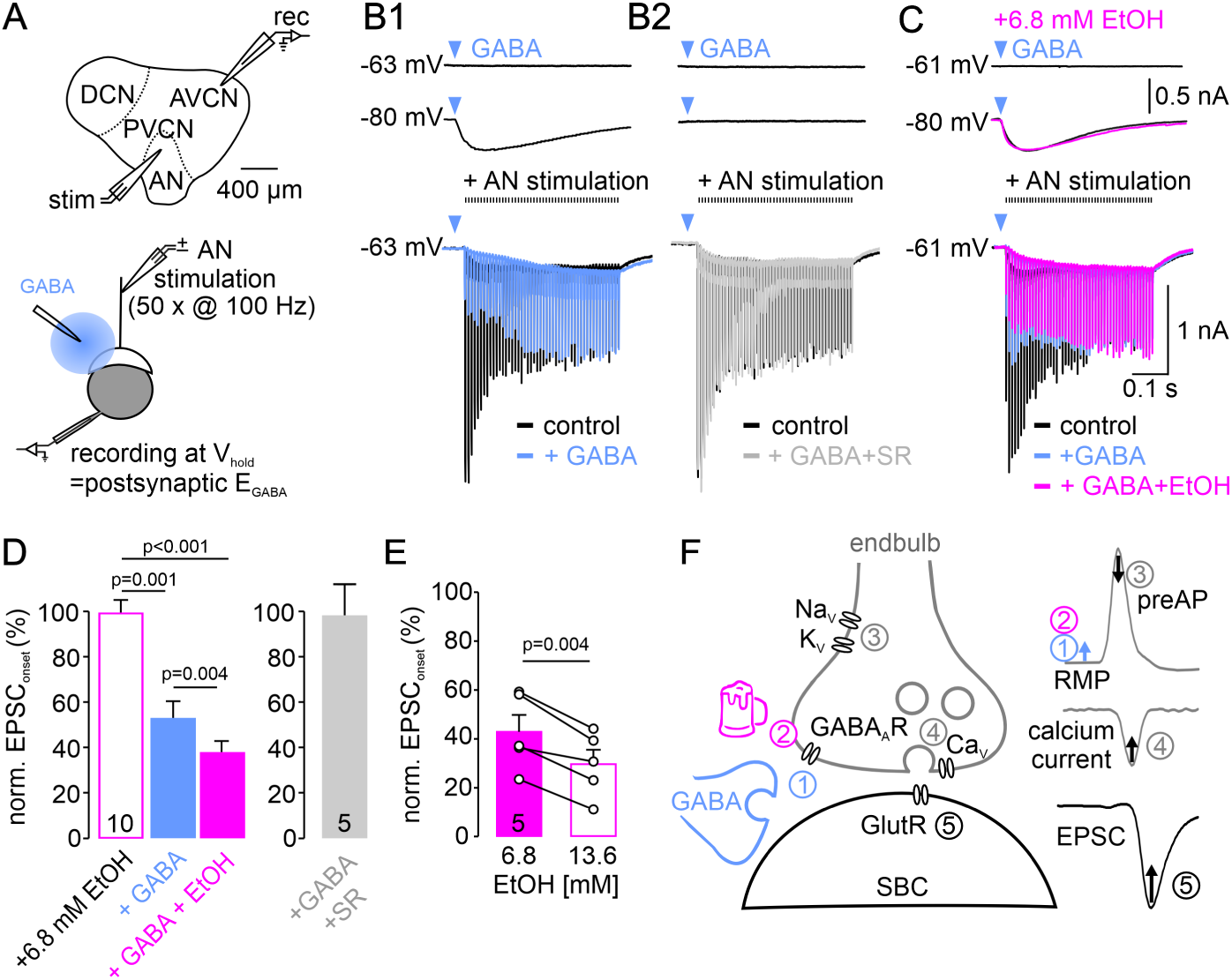
Ethanol potentiates presynaptic GABA_A_R-mediated suppression of synaptic transmission. (A) Schematic of the *in vitro* recording configuration. EPSCs were evoked by auditory nerve (AN) stimulation while voltage-clamping the postsynaptic SBC at its GABA reversal potential (V_hold_ = E_GABA_) to eliminate postsynaptic GABAergic currents. GABA was locally applied to the auditory nerve terminal (bottom). (B) Isolation of presynaptic inhibition. Top: Responses to local GABA puffs showing inward current at V_hold_ = −80 mV and no net current at E_GABA_. Bottom: High-frequency EPSC trains (100 Hz) recorded at E_GABA_. (B1) Local GABA application (blue) suppresses EPSC amplitudes. (B2) The GABA_A_R antagonist SR95531 abolishes this suppression. (C) Ethanol enhances synaptic suppression. Bath application of ethanol (6.8 mM, magenta) further reduced EPSC amplitudes compared to GABA alone (blue). Note that postsynaptic GABA currents (top traces) remain unchanged in the presence of ethanol, confirming a presynaptic site of action. (D) Quantification of normalized EPSC onset amplitudes (pulses 1-5). Ethanol significantly potentiates the GABA-mediated suppression of release. (E) Dose-dependent effect. Doubling the ethanol concentration (13.6 mM) further suppressed synaptic transmission. (F) Proposed mechanism: (1) GABA activates presynaptic GABA_A_R, causing (2) terminal depolarization, which is potentiated by ethanol. This depolarization (3) recruits Kv1 channels to shunt the presynaptic AP, thereby (4) reducing Ca^2+^ influx and (5) suppressing glutamate release.

Application of GABA (500 µM, 10 ms) reduced EPSC amplitudes by 46 ± 7 % (mean of EPSCs 1-5), an effect blocked by SR95531 (Fig. 7B, D; EPSC: control = 2.0 ± 0.7 nA, GABA+SR = 1.9 ± 0.7 nA, control vs. SR95531 p = 0.59, n=5, paired t-test). Bath application of 6.8 mM ethanol significantly enhanced this suppression to 63 ± 5% (Fig. 7C, D; EPSC: control = 2.4 ± 0.5 nA, GABA = 1.3 ± 0.4 nA, GABA+EtOH = 1.0 ± 0.3 nA, control vs. GABA p = 0.003, GABA vs. GABA+EtOH p = 0.004, n=10, one-way RM ANOVA). The ethanol-induced suppression closely followed the time course of the presynaptic GABA_A_R current (Fig. 7C cf. Fig. 2J; I_GABA_: 10-90% rise = 50.9 ± 7.5 ms, 90-10% decay = 423.6 ± 64.6 ms, n=12). Ethanol alone had no effect on EPSC amplitudes (Fig. 7D; control vs. +EtOH p = 0.95, one-way RM ANOVA) or on postsynaptic GABA currents (Fig. 7C), confirming that its action requires presynaptic GABA_A_R activation. Doubling the ethanol concentration to 13.6 mM (∼0.6 ‰) further reduced EPSCs to 30 ± 6 % of control (Fig. 7E; EPSC: 6.8 mM = 0.8 ± 0.2 nA, % of control = 43 ± 7 %; 13.6 mM = 0.6 ± 0.1 nA, 6.8 mM vs. 13.6 mM p = 0.034, n=5, one-way RM ANOVA). These data show that ethanol dose-dependently potentiates GABA_A_R-dependent suppression of synaptic transmission under physiologically relevant firing conditions.

## Discussion

Our data reveal that ethanol suppresses sound-evoked activity in SBCs via GABA_A_R *in vivo* (Fig. 1) and that potentiation of presynaptic GABA_A_R provides a cellular mechanism for ethanol-induced auditory impairment (Fig. 7F). We show that ethanol concentrations approximating those reached after a single alcoholic beverage selectively enhance a depolarizing chloride conductance at the endbulb of Held terminal, depolarizing it from its resting membrane potential (RMP, cf. ① and ② in Fig. 7F). This depolarization opens low-voltage-activated Kv1 channels, which shunt presynaptic APs and reduce their amplitude (③), in turn suppressing calcium influx (④) and glutamate release (⑤). These findings identify a synaptic mechanism linking acute alcohol consumption to disrupted auditory processing.

### Ethanol suppresses auditory gain through GABAergic inhibition

Our *in vivo* recordings showed that behaviorally relevant ethanol concentrations suppress sound-evoked firing in SBCs, particularly at sound intensities relevant for speech communication, without affecting response thresholds or temporal precision. This suppression was reversed by the GABA_A_R antagonist SR95531, implicating GABAergic inhibition as the primary mediator. The preservation of sensitivity and temporal fidelity despite reduced synaptic gain narrows the number of potential mechanisms. In the auditory brainstem, inhibition is predominantly glycinergic and postsynaptic (Moore and Caspary, 1983; Wu and Kelly, 1992; Kandler and Friauf, 1995; Golding and Oertel, 1996; Awatramani et al., 2004; Pecka et al., 2008; Nerlich et al., 2014a), operating via a subtractive mechanism that filters weak inputs and sharpens temporal precision (Myoga et al., 2014; Franken et al., 2015; Keine et al., 2016; Beiderbeck et al., 2018). In contrast, ethanol-induced suppression points to modulation of presynaptic efficacy rather than postsynaptic excitability. Consistent with this, ethanol had no effect on postsynaptic GABA_A_R conductance at the endbulb of Held – SBC synapse.

### Presynaptic GABA_A_R are the primary ethanol target at the endbulb synapse

Ethanol can modulate a broad range of ion channels, including GlyR (Aguayo and Pancetti, 1994; Aguayo et al., 1996; Yevenes et al., 2008; Welsh et al., 2009; Burgos et al., 2015), BK channels (Dopico et al., 2016), nAChR (Covernton and Connolly, 1997), NMDAR (Lovinger et al., 1990), AMPAR (Martin et al., 1995), and P2XR (Ostrovskaya et al., 2011). Within this landscape, our data identify presynaptic GABA_A_R as the principal ethanol target at the first central auditory synapse. GABA_A_R on SBCs were insensitive to ethanol, whereas receptors at the endbulb exhibited robust potentiation. This differential sensitivity likely reflects differences in subunit composition. The dose-dependent potentiation by behaviorally relevant ethanol concentrations (6.8 vs. 13.6 mM) and the lack of postsynaptic effects are characteristic of high-affinity, extrasynaptic GABA_A_R (Lobo and Harris, 2008). Two subunit features have been implicated as determinants of low-dose ethanol sensitivity: the β3 subunit (Wallner et al., 2003) and the δ subunit, which endows receptors with high GABA affinity and extrasynaptic localization (Brickley and Mody, 2012). Several studies have reported that δ containing GABA_A_R are potentiated by low ethanol concentrations (Sundstrom-Poromaa et al., 2002; Wallner et al., 2003; Wei et al., 2004; Nie et al., 2011), yet this finding has not been universally reproduced (Borghese et al., 2006; Yamashita et al., 2006; Casagrande et al., 2007; Korpi et al., 2007). Such discrepancies may in part reflect variable δ-subunit incorporation in recombinant expression systems (Meera et al., 2010) and differences in intracellular signaling context, as ethanol potentiation of δ-containing GABA_A_R requires protein kinase Cδ activity (Choi et al., 2008).

### Depolarizing chloride efflux underlies presynaptic GABA_A_R action

Presynaptic inhibition via metabotropic GPCRs such as GABA_B_R is a ubiquitous feature of mammalian central synapses (White and Tyler, 1987; Dittman and Regehr, 1996; Wu and Saggau, 1997; Bettler et al., 2004; Chalifoux and Carter, 2011; Laviv et al., 2011) with the notable exception of the medial habenula–interpeduncular nucleus synapse, where GABA_B_R activation potentiates release (Koppensteiner et al., 2024). GABA_A_R-mediated presynaptic inhibition, although less common, is prominent in the mammalian spinal cord (Eccles et al., 1963; Nicoll and Alger, 1979), where it underlies smooth movement (Fink et al., 2014). Presynaptic GABA_A_R-dependent modulation has also been shown in the retina (Tachibana and Kaneko, 1987; Lukasiewicz and Werblin, 1994), posterior pituitary (Saridaki et al., 1989; Zhang and Jackson, 1993), and auditory brainstem (Lim et al., 2000; Turecek and Trussell, 2002). The functional consequence of presynaptic GABA_A_R activation is synapse-specific: in the hippocampus, these receptors facilitate release at both mossy fiber boutons (Ruiz et al., 2010) and Schaffer collateral terminals (Jang et al., 2005), whereas at cerebellar parallel fibers the effect depends on the level of receptor activation (Khatri et al., 2019), and at other CNS terminals presynaptic GABA_A_R can suppress release (Wakita et al., 2012).

The mechanisms of GABA_A_R-mediated presynaptic modulation are incompletely understood. A central finding of the present study is that GABA depolarizes the endbulb of Held terminal due to an outwardly directed chloride gradient (Fig. 3). Our results thus provide experimental support for the long-standing model of presynaptic inhibition in the dorsal roots of the spinal cord, in which GABA_A_R-mediated depolarization is measured as “primary afferent depolarization” (Eccles et al., 1963; Barber et al., 1978; Wall and Lidierth, 1997; Witschi et al., 2011). At the calyx of Held synapse in the brainstem, glycine induces a similar presynaptic depolarizing chloride gradient (Price and Trussell, 2006). However, at the calyx and several central synapses, presynaptic depolarization facilitate neurotransmitter release either by broadening the action potential (Alle and Geiger, 2006; Shu et al., 2006; Kole et al., 2007) or by elevating basal Ca^2+^ (Turecek and Trussell, 2001; Awatramani et al., 2005; Alle and Geiger, 2006; Shu et al., 2006; Eshra et al., 2021). In contrast, our data demonstrate that a depolarization at the endbulb of Held terminal suppresses glutamate release (Fig. 4, 6, 7). A related depolarizing shunt has been described postsynaptically in the chick nucleus laminaris, where δ-subunit-containing extrasynaptic GABA_A_R generate a tonic depolarizing conductance that sharpens coincidence detection (Tang et al., 2011). Our findings extend this depolarizing shunting principle to a presynaptic auditory terminal.

### Recruitment of Kv1 channels converts depolarization into AP shunting

Presynaptic Kv1 channels appear to play a distinctive role at the endbulb – SBC synapse. While depolarization inactivates Kv1 channels in the hippocampus and cortex, thereby broadening the AP (Alle and Geiger, 2006; Shu et al., 2006; Kole et al., 2007), Kv1 channels at the endbulb remain available at GABA-induced membrane potentials and are further recruited, leading to AP attenuation rather than broadening (Fig. 5). Our DTX-K experiments show that Kv1 channels at the endbulb share the low activation threshold and minimal inactivation previously described in bushy cells (Rothman and Manis, 2003; Cao et al., 2007), MNTB principal neurons (Brew and Forsythe, 1995) and at the calyx of Held presynaptic terminal (Dodson et al., 2003; Ishikawa et al., 2003). Consequently, the GABA-induced reduction in AP amplitude translates into a pronounced reduction of the EPSC, reflecting the supralinear relationship between presynaptic calcium influx and synaptic vesicle fusion (Schneggenburger and Neher, 2000; Sakaba and Neher, 2001; Wang et al., 2008; Lin et al., 2011). Thus, by potentiating presynaptic GABA_A_R, ethanol amplifies the chloride conductance, further recruiting Kv1 channels and exacerbating suppression of glutamate release. This presynaptic suppression of AP amplitudes without temporal broadening translates directly into compromised intensity coding with preserved temporal precision, as observed in our *in vivo* recordings.

### Possible sources of GABA

If ethanol acts via presynaptic GABA_A_R, what is the source of their GABA ligand? SBCs receive acoustically evoked inhibition that enhances temporal precision relative to the auditory nerve input and improves the reproducibility of sound evoked responses (Dehmel et al., 2010; Kuenzel et al., 2011; Keine et al., 2016, 2017). This primarily glycinergic inhibition is amplified by co-release with GABA, creating a dynamic high-pass filter (Nerlich et al., 2014b; Nerlich et al., 2017). Due to slow transmitter clearance and asynchronous release, this inhibitory conductance builds up at physiological firing rates, providing a tonic-like inhibition (Nerlich et al., 2014a). *In vivo*, inhibition onto SBCs has a higher activation threshold than excitation, and therefore preferentially suppresses firing at elevated sound levels (Kuenzel et al., 2011; Keine and Rübsamen, 2015; Kuenzel et al., 2015; Keine et al., 2016). This matches the intensity profile of ethanol-induced firing suppression we observed at higher intensities, likely driven by GABA spillover onto the presynaptic terminal. Indeed, GABAergic terminals lacking direct postsynaptic receptor targets have been identified near SBCs (Nerlich et al., 2017), raising the possibility that such terminals activate presynaptic GABA_A_R via volume transmission. Such spillover-mediated activation may provide a mechanism for activity-dependent control of synaptic strength at the endbulb via GABA_A_Rs, complementing the previously proposed regulation through GABA_B_Rs (Chanda and Xu-Friedman, 2010). Notably, extrasynaptic GABA_A_R may sense GABA primarily as a steady-state ambient signal rather than as discrete spillover transients, because high-affinity δ-containing receptors desensitize rapidly in the presence of low ambient GABA (Bright et al., 2011). The activity-dependent build-up of inhibition at SBCs (Nerlich et al., 2014a) is well suited to provide such a tonic ambient signal.

### A synaptic basis for ethanol-induced auditory deficits

The gain reduction observed at the first central auditory synapse plausibly compromises the detection of signals in noise, where rate-based coding of sound envelopes is required to represent signals against a fluctuating background (Frisina, 2001; Joris et al., 2004). Critically, AP attenuation at the endbulb occurs without temporal broadening, preserving presynaptic spike timing and downstream temporal precision, consistent with the unchanged vector strength observed under ethanol. Instead, rate-level functions were flattened, likely reducing the dynamic range over which sound intensity is encoded. This selective degradation of rate-based coding at the central auditory relay provides a plausible cellular mechanism for auditory deficits in which speech discrimination in noise is impaired even at low blood-alcohol levels (Upile et al., 2007; Choi et al., 2021), despite preserved pure-tone sensitivity and temporal fidelity.

Previous work has established that low ethanol concentrations produce only modest reductions in cochlear gain mediated by outer hair cells (Hwang et al., 2003; Liu et al., 2004), consistent with minimal shifts in pure-tone thresholds observed in humans and rodents (Murata et al., 2001; Hwang et al., 2003; Liu et al., 2004; Choi et al., 2021). Our findings reveal a second stage of ethanol-induced auditory dysfunction: a substantially larger reduction in synaptic gain at the endbulb of Held – SBC synapse (Fig. 1, 7), the consequences of which propagate through the ascending pathway. The preserved temporal precision in our recordings aligns with reports of intact gap-in-noise detection, word recognition in quiet, and selective listening among competing speakers (Choi et al., 2021; Harvey and Beaman, 2021). The robust suppression of firing rates we observed (Fig. 1), by contrast, plausibly underlies the impairment of speech-in-noise perception reported in behavioral studies (Choi et al., 2021) and the attenuated cortical responses to sound changes at comparable blood-alcohol levels (Kähkönen et al., 2005; He et al., 2013). We therefore propose that ethanol-induced reduction of synaptic gain at the first central synapse degrades the signal-to-noise ratio available for downstream circuits required for auditory stream segregation, thereby contributing to “cocktail-party deafness”. A brainstem origin of this impairment is consistent with the observation that ethanol prolongs auditory brainstem response latencies beginning at the generator associated with the cochlear nucleus (Chu et al., 1978; Church and Williams, 1982) and that cortical event-related potentials are correspondingly suppressed (Jääskeläinen et al., 1996; He et al., 2013). Together, these findings establish presynaptic shunting inhibition at the first central auditory synapse as a cellular substrate for ethanol-induced auditory deficits, identifying a synaptic mechanism by which acute, low-dose ethanol degrades signal representation at the earliest stage of central auditory processing.

## Methods

### Animals

All experiments were conducted at the Physiology Laboratory or the General Zoology and Neurobiology Laboratory at the University of Leipzig in accordance with the German Animal Welfare Act (§4 TSchG) and the European Union Directive 2010/63/EU, under protocols approved by the Saxony State Animal Welfare Committee (TVV 06/09). We used Mongolian gerbils (*Meriones unguiculatus*) of either sex, aged postnatal day (P) 13–P31, group-housed under a standard 12-h light/dark cycle with *ad libitum* access to food and water. Every effort was made to minimize the number of animals used and to alleviate pain or distress.

### *In vivo* electrophysiology

#### Surgical preparation

Recordings were obtained from 9 gerbils (P24-31) of either sex (5 females). Anaesthesia was induced via intraperitoneal injection of ketamine hydrochloride (140 mg/kg body weight; Ketamin-Ratiopharm, Ratiopharm) and xylazine hydrochloride (3 mg/kg body weight; Rompun, Bayer) and maintained by subcutaneous injections (one-third of the initial dose) approximately every 90 min. Body temperature was maintained between 37.5 and 38.5°C using a feedback-controlled heating pad.

Following exposure of the skull along the dorsal midline, a metal bolt was glued to bregma to fix the animal in the stereotaxic frame within a sound-attenuated chamber (Type 400, Industrial Acoustic Company). Two craniotomies were performed in the occipital bone, 1.8–2.3 mm caudal to lambda: a medial hole (Ø 0.5 mm) for the reference electrode, placed in the superficial cerebellum, and a lateral hole (Ø 1 mm, 1 mm lateral to midline) for the recording electrode. To target the rostral, low-frequency pole of the anteroventral cochlear nucleus (AVCN), the head was tilted 6–10° rostro-caudally and 14–16° mediolaterally.

#### Single-unit recording

Acoustic stimuli were digitally synthesized (Matlab, The MathWorks Inc.) and delivered via a D/A converter (RP2.1 real-time processor, 97.7 kHz sampling rate; Tucker-Davis Technologies) to custom earphones (acoustic transducer: DT 770 Pro, Beyer Dynamics) fitted with plastic tubing (length: 35 mm; inner diameter: 5 mm) inserted into the outer ear canal.

We located the rostral AVCN (characteristic frequency, CF = 2.6±1.9 kHz, n=11) by monitoring multi-unit activity to low-frequency sounds with low-impedance glass micropipettes (GB150F-10, Science Products; 1–5 MΩ, filled with 3 M KCl). Juxtacellular single-unit recordings were obtained using three- or four-barrelled piggyback electrodes (tip diameter: 4–6 µm; recording barrel protrusion: 15–25 µm; impedance: 5–10 MΩ; GB200F-10, 3GB120F-10, 4GB120F-10, Science Products) as described previously (Havey and Caspary, 1980; Dietz et al., 2012; Keine et al., 2016).

Excitatory response areas were mapped using pure tone pulses (100 ms duration, 5 ms rise–fall time, 100 ms interstimulus interval), presented in a frequency-intensity matrix (20 frequencies on a logarithmic scale, 10 intensities on a linear scale, 4-5 repetitions). The CF was defined as the frequency eliciting a response above spontaneous firing rate at the lowest stimulus intensity (i.e., the tip of the excitatory tuning curve).

Pharmacological agents were applied iontophoretically (EPMS 07, npi electronics). To prevent passive leak, retention currents of 20 nA (opposite polarity of ejection) were applied between drug deliveries. Ethanol (40 mM, pH 8) and SR95531 (5 mM, pH 4) were ejected using currents ranging from −50 to −150 nA and +25 to +100 nA, respectively. A balancing barrel (1M sodium acetate) was used to offset capacitive artifacts. Control applications of the vehicle alone (distilled water, pH 8) at −50 to −150 nA produced no effect. Spontaneous activity was assessed in the absence of acoustic stimulation, before, during, and after drug application.

### *In vitro* electrophysiology

#### Acute slice preparation

Animals (P13–P30, n=115, 61 female) were deeply anesthetized with isoflurane (Baxter Deerfield, IL) and decapitated. The brain was extracted and parasagittal slices (200 µm) containing the cochlear nucleus were prepared using a vibratome (Leica VT1200 S) in ice-cold (3–4°C) low-calcium, sucrose-based solution (in mM): 215 sucrose, 2.5 KCl, 0.1 CaCl_2_, 4 MgCl_2_, 1.25 NaH_2_PO_4_, 25 NaHCO_3_, 10 glucose, 2 sodium pyruvate, 3 myo-inositol, and 0.5 ascorbic acid. Slices were incubated for 30 min at 37°C in artificial cerebrospinal fluid (ACSF) containing (in mM): 125 NaCl, 2.5 KCl, 1.2 CaCl_2_, 1 MgCl_2_, 1.25 NaH_2_PO_4_, 25 NaHCO_3_, 10 glucose, 2 sodium pyruvate, 3 myo-inositol, 0.5 ascorbic acid (equilibrated with carbogen, pH 7.4) and stored at room temperature until recording.

#### General recording conditions

Unless otherwise noted, recordings were performed at near-physiological temperature (34 ± 0.5°C) in ACSF supplemented with antagonists for GABA_B_ receptors (CGP 55845, 3 µM), NMDA receptors (D-AP5, 50 µM) and glycine receptors (strychnine, 0.5 µM) to pharmacologically isolate GABA_A_-mediated and AMPA-mediated transmission. Signals were acquired using a Multiclamp 700B amplifier (Molecular Devices) and low-pass filtered at 6 kHz (4-pole Bessel, voltage-clamp) or 10 kHz (current-clamp) and digitized at 20 kHz (voltage-clamp) or 100 kHz (current-clamp).

Agonists (glycine, GABA, muscimol) were pressure-applied via micropipettes (3 µm tip, 3 psi, 10 ms duration, Picospritzer, General Valve Corp) positioned ∼10 µm from the target structure.

Ethanol (Rotipuran®, ≥99.8 %, analytical grade; Carl Roth, Germany) was bath-applied at concentrations of 6.8 mM or 13.6 mM.

In a subset of experiments, GABA induced depolarizations were mimicked by 10 mV ramped depolarizations (70 ms duration) via current injection (Fig. 4E).

##### Postsynaptic whole-cell recordings from SBCs

Whole-cell recordings from SBCs in the AVCN were obtained using borosilicate glass pipettes filled with (in mM): 130 CsMeSO_3_, 10 TEA-Cl, 10 HEPES, 5 EGTA, 3 Mg-ATP, 0.3 Na_2_-GTP, 5 Na_2_-phosphocreatine and 5 QX314-Cl (pH 7.3, 298 mOsm). Series resistance (11.7 ± 0.5 MΩ, n=84) was compensated by 50–70%. To isolate EPSCs, the holding potential was set to the experimentally determined equilibrium potential of GABA_A_R (E_GABA_) for each cell (range – 58 to –69 mV, see Fig. S4). All voltages were corrected offline for a calculated liquid junction potential of 10 mV. EPSCs were evoked by electrical stimulation of the auditory nerve root using a tungsten concentric bipolar electrode (World Precision Instruments) driven by an isolated stimulus unit (Iso-flex, AMPI). Stimulus trains consisted of 50 pulses at 100 Hz, delivered every 15 s. To assess the effects on glutamatergic transmission, GABA (100, 500 µM), glycine (500 µM) or muscimol (100 µM) were pressure applied during synaptic stimulation.

##### Presynaptic whole-cell recordings from endbulbs of Held

Endbulb terminals were visually identified under differential inference contrast (DIC) optics and targeted using pipettes containing ATTO 488 (25 µM, ATTO-TEC, Germany) for confirmation of the recording site.

Voltage-clamp recordings: Pipettes were filled with (in mM): 120 CsCl, 20 TEA-Cl, 1 MgCl_2_, 20 HEPES, 5 EGTA, 2 Na_2_-ATP, 0.3 Na_2_-GTP, 5 Na_2_-phosphocreatine and 5 QX314-Cl (pH 7.3, 298 mOsm). The holding potential was set to −72 mV after correction for a 2 mV liquid junction potential. Series resistance was 17.3 ± 1.07 MΩ (mean ± SEM, n=50) and was not compensated due to relatively small voltage errors.

Current-clamp recordings: Pipettes contained (in mM): 100 K-MeSO_3_, 40 KCl, 10 HEPES, 0.1 EGTA, 3 Mg-ATP, 0.3 Na_2_-GTP, 5 Na_2_-phosphocreatine (pH 7.3, 298 mOsm). Voltages were corrected for a calculated 8.6 mV liquid junction potential and bridge balance was adjusted to 100% of the series resistance. APs were evoked either by electrical stimulation of the auditory nerve root (see above) or by suprathreshold current injection into the terminal (0.1 ms square pulses). Stimulation patterns included single pulses at 0.1 Hz and spike trains derived from *in vivo* recordings of SBC spontaneous and sound-evoked activity (Fig. 4).

##### Presynaptic perforated-patch recordings

To preserve native intraterminal chloride concentrations, gramicidin-perforated patch recordings were performed as previously described (Nerlich et al., 2014b). These experiments were conducted at 28 ± 0.5 °C to optimize seal stability during the prolonged perforation period. Pipettes were pulled from borosilicate glass to a resistance of 3.5-4.5 MΩ and tip-filled with (in mM): 147 K-gluconate, 5 KCl, 1 MgCl_2_, 10 HEPES, 5 EGTA (pH 7.3, 303 mOsm). The remainder of the pipette was back-filled with the same solution supplemented with gramicidin A (30 µg/ml, Sigma) and ATTO 488 (25 µM) to verify patch integrity via fluorescence monitoring. Perforation was monitored by the increase in steady-state current in response to a −5 mV voltage step. Series resistance typically stabilized within 5-20 min (mean ± SEM = 11.9 ± 1.4 min, n=16) at 32.6 ± 2.5 MΩ (n=16) and was not compensated. E_GABA_ was determined by linear regression of the current-voltage relationship of GABA-evoked currents. To assess the contribution of HCO_3_ ^−^ to the E, a subset of recordings was performed in HEPES-buffered ACSF (NaHCO_3_ replaced by 10 mM HEPES). Perforated patch integrity was validated in separate control experiments using a high-chloride pipette solution (K-gluconate replaced by KCl; [Cl^−^]_pip_ = 150 mM). All voltages were corrected for liquid junction potentials of 1.4 mV ([Cl^−^]_pip_=150 mM) or 13.2 mV ([Cl^−^]_pip_=7 mM).

##### Simultaneous pre- and postsynaptic recordings

To relate presynaptic calcium influx to neurotransmitter release, simultaneous whole-cell recordings were established from synaptically connected endbulb of Held terminals and postsynaptic SBCs. Presynaptic calcium currents were pharmacologically isolated by reducing the external NaCl concentration to 105 mM and supplementing ACSF with (in mM): 10 TEA-Cl, 2 4-AP, 0.001 TTX, 0.05 ZD 7288, 0.02 SR95531. The presynaptic pipette solution contained (in mM): 105 CsMeSO_3_, 20 CsCl, 20 TEA-Cl, 10 HEPES, 0.1 EGTA, 2 Mg-ATP, 0.3 Na_2_-GTP, 5 Na_2_-phosphocreatine and 3 Na-L-glutamate (pH 7.31, 294 mOsm). The postsynaptic pipette solution was identical to that used for isolated SBC recordings (see above). The endbulb was held at −77 mV and the SBC at −80 mV, after correcting for calculated junction potentials of 9.3 mV (presynaptic) and 10 mV (postsynaptic). To mimic physiological activation, presynaptic calcium currents and resulting EPSCs were evoked using voltage-command waveforms derived from averaged APs recorded during auditory nerve stimulation (see above). In a subset of experiments designed to assess voltage-dependent calcium channel inactivation, calcium currents were elicited by a 0.3 ms step to 0 mV, preceded by depolarizing ramps of increasing amplitude (5 mV increments, 70 ms, Fig. 6).

### Data analysis

#### In vivo data analysis

Signals were amplified (Neuroprobe 1600, A-M Systems), bandpass-filtered (0.3-7 kHz), and digitized at 97.7 kHz (RP2.1, Tucker-Davis Technologies). Single-units were isolated using custom Matlab functions (B. Englitz, Department of Neurophysiology, Nijmegen) based on a signal-to-noise ratio >8:1 and uniform waveforms. SBCs were identified by their characteristic complex waveform, comprising the presynaptic endbulb potential (“prepotential”) and a postsynaptic AP (Pfeiffer, 1966; Young et al., 1988; Englitz et al., 2009; Typlt et al., 2010; Typlt et al., 2012), and a primary-like peristimulus time histogram (Blackburn and Sachs, 1989). Stellate cells were excluded based on their biphasic waveforms and ‘chopper’ responses (Rhode and Smith, 1986; Young et al., 1988; Typlt et al., 2012).

#### In vitro data analysis

##### Calculation of presynaptic E_GABA_ and chloride concentration

E_GABA_ was calculated from gramicidin-perforated patch recordings and corrected for the voltage drop across the series resistance. The physiological presynaptic [Cl]_i_ was calculated using the Nernst equation [Cl]_i_ = [Cl]_o_ *e*^EGABA^*F/RT*, where [Cl]_o_ represents the extracellular chloride concentration (132 mM), E_GABA_ is the reversal potential of the GABA-evoked current, *F* is the Faraday constant, *R* is the gas constant, and *T* is the temperature (28°C / 301 K).

##### EPSC analysis

EPSC peak amplitudes were quantified from averaged traces (>5 repetitions) using custom-written Matlab routines. To characterize synaptic dynamics during high-frequency stimulation (50 pulses at 100 Hz), we defined “onset” amplitude as the mean of the first 5 EPSCs and “steady state” amplitude as the mean of the 17th to 26th EPSCs. In pharmacological experiments, agonists were pressure-applied 10ms prior to the onset or steady-state analysis window.

##### Presynaptic APs

AP amplitude was calculated as the voltage difference between the pre-stimulation baseline and the AP peak, and AP half-width as the duration at half-maximal amplitude. Measured AP amplitudes were smaller than those reported in mouse endbulbs (Lin et al., 2011), which could potentially result from capacitive filtering due to uncompensated pipette capacitance (Oláh et al., 2021; Ritzau-Jost et al., 2021). However, several factors argue against such a major technical limitation in our recordings: (i) the relatively large capacitance of gerbil endbulbs (∼9 pF) compared to small boutons (∼1 pF) reduces susceptibility to filtering; (ii) pipette capacitance was minimized, and recordings were performed with a current-clamp amplifier optimized to reduce capacitive filtering (MultiClamp 700A; Oláh et al., 2021; Ritzau-Jost et al., 2021); and (iii) pharmacological broadening of APs with TEA increased AP half-width without affecting amplitude (Fig.5A-C), which argues against a dominant contribution of capacitive filtering.

### Statistics

Individual SBCs were treated as independent biological samples. Sample sizes were chosen based on prior studies of similar synaptic recordings (Nerlich et al., 2014a; Nerlich et al., 2014b). Aggregated data are presented as mean ± SEM, unless otherwise noted. All data sets were tested for normal distribution with Shapiro-Wilk tests. Pairwise comparisons were performed using paired t-test for within-subject design and unpaired Student t-test for independent groups. Comparisons involving more than two groups were assessed by one-way ANOVA, while multifactorial designs were analyzed using two-way ANOVA, followed by Holm-Sidak post-hoc tests. Statistical analyses were conducted using Sigma Plot (Version 11.0, Systat Software), Matlab or Jamovi (https://www.jamovi.org/).

