## Supplemental Figures for "Presynaptic GABA_A_ Receptors Mediate Ethanol-Induced Suppression at a Central Auditory Synapse"

### Supplemental data

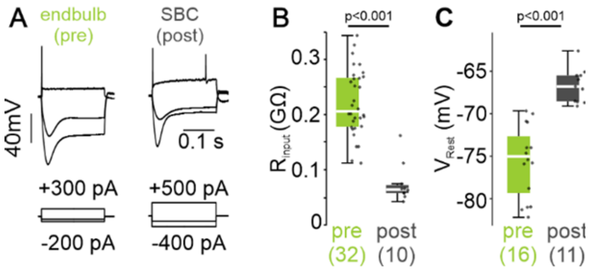


**Fig. S1 Intrinsic properties of pre- and postsynaptic elements**

(A) Voltage responses of the endbulb of Held and SBC to hyperpolarizing and depolarizing current injections, showing single AP firing in response to depolarization.

(B-C) Input resistance (B) and resting membrane potential (C) for endbulbs (pre) and SBCs (post). Box plots show the median and interquartile range; whiskers extend to 1.5 x IQR. Individual points represent single cells, with sample sizes indicated in parentheses. *p* values were obtained using unpaired t-tests.

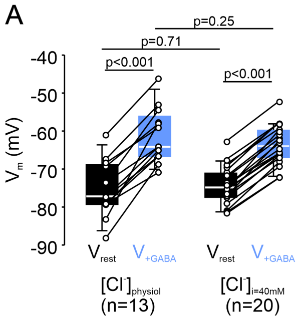


**Fig. S2 Comparable membrane potentials with physiological intraterminal [Cl$^-$]$_i$ and [Cl$^-$]$_i$ = 40 mM**

(A) Resting membrane potential ($V_{rest}$) and membrane potential during GABA application ($V_{+GABA}$) for recording with physiological [Cl$^-$]$_i$ (gramicidin-perforated patch, n=13) and with [Cl$^-$]$_i$ = 40 mM (whole-cell, n=20). Box plots show the median and interquartile range; *p* values were obtained using two-way ANOVA.

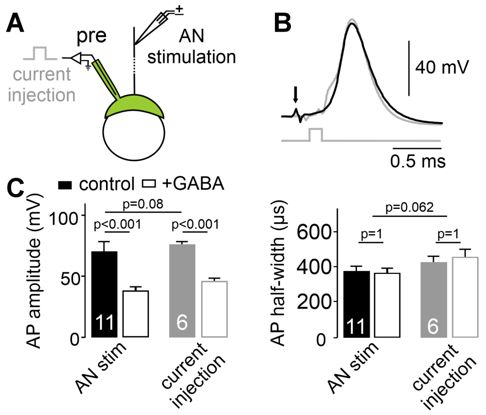


**Fig. S3 Action potential generation paradigm does not influence AP properties**

(A) Presynaptic recording configuration. APs in the endbulb were evoked either by auditory nerve (AN) stimulation or by brief depolarizing current injection (0.1 ms).

(B) Overlaid example traces of APs evoked by AN stimulation (black) or current injection (grey), showing similar waveforms.

(C) Mean AP amplitudes (left) and AP half-widths (right) under control condition and during presynaptic GABA application. GABA significantly reduced AP amplitude in both paradigms, whereas AP half-width was unchanged. *p* values were obtained using two-way ANOVA. Cell numbers are shown within the bars.

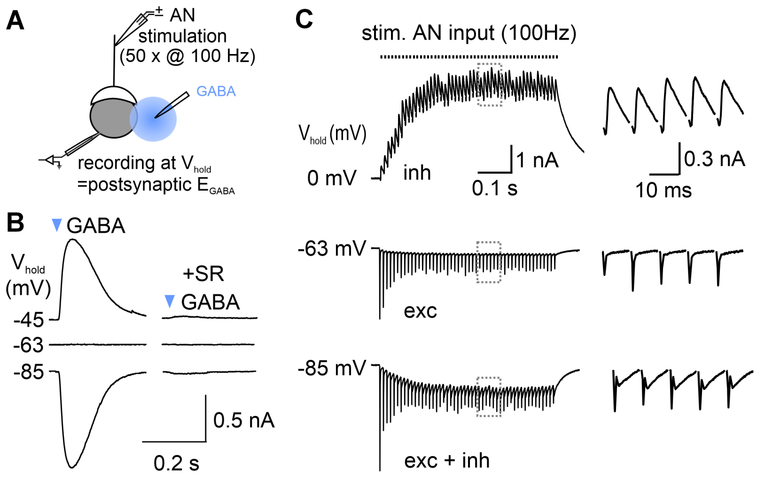


**Fig. S4 Isolation of EPSCs at $V_{hold}$ = $E_{GABA}$**

(A) Postsynaptic recording configuration with auditory nerve stimulation (used in C) and local GABA application near the SBC soma (used in B).

(B) GABA-evoked currents recorded at different holding potentials to determine postsynaptic $E_{GABA}$; currents are abolished by SR95531, confirming $GABA_AR$ specificity.

(C) Postsynaptic responses to AN stimulation at different holding potentials. At $V_{hold}$ = 0 mV ($E_{AMPA}$), isolated inhibitory postsynaptic potentials (IPSCs) show slow decay; at -63 mV ($E_{GABA}$), isolated EPSCs show fast decay; at

-85 mV, mixed EPSCs and IPSCs are observed. Insets show overlaid EPSC and IPSC traces, highlighting delayed IPSC onset consistent with a disynaptic origin.

Several control experiments were performed to substantiate the role of $GABA_AR$ in modulating glutamatergic transmission.

Delayed GABA application during the steady state phase of the EPSC train (mean of EPSC 17 to 26) reduced EPSC amplitudes by 49 ± 7%, and this effect was blocked by SR95531 in a subset of experiments (Fig. S5B and C). We also confirmed that this presynaptic GABAergic modulation persists into later developmental stages (P25-30), with EPSC reduction still evident (Fig. S5A and C, EPSC reduction P25-30: onset = 46 ± 9 %, p = 0.002, steady state = 37 ± 8 %, n=8, p = 0.005, paired t-tests).

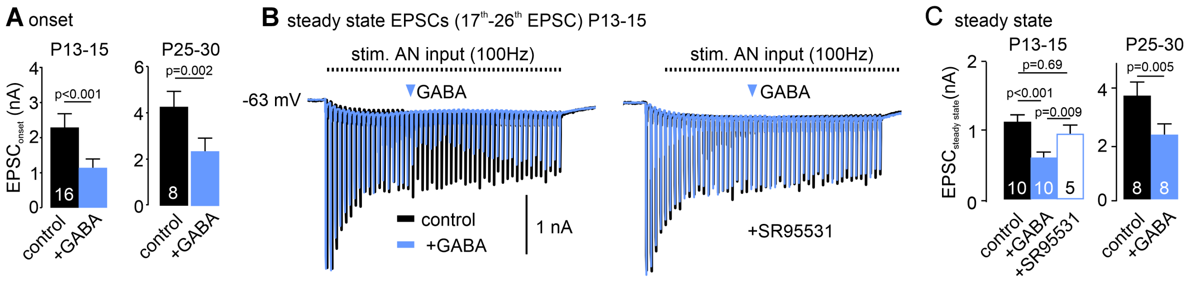


**Fig. S5 Presynaptic $GABA_AR$ reduce onset and steady-state EPSC amplitudes throughout development**

(A) Onset EPSC amplitudes (mean of EPSCs 1-5) are reduced by 500 µM GABA in both juvenile (P13-15; n=16, paired t-test) and young adult gerbils (P25-P30; n=8, paired t-test).

(B) Example EPSCs recorded at $V_{hold}$ = $E_{GABA}$ before and during GABA application; the steady-state reduction is abolished by SR95531.

(C) Steady-state EPSC amplitudes (mean of EPSCs 17-26) are likewise reduced by GABA at both ages (P13–15, RM ANOVA, n=5; P25–30, paired t-test). Numbers within the bars indicate sample sizes.

Application of the specific $GABA_AR$ agonist muscimol (100 µM) reduced EPSCs amplitudes by 32 ± 3 % at train onset and by 29 ± 6% during the steady state (Fig. S6A and B, $EPSC_{onset}$: control = 2.7 ± 0.6 nA, mucimol = 1.8 ± 0.4 nA, $EPSC_{steady}$: control = 2.2 ± 0.6 nA, mucimol = 1.5 ± 0.4 nA, n=5, control vs. muscimol p = 0.013, two-way RM ANOVA). This further supports a $GABA_AR$-dependent mechanism of suppression of glutamatergic transmission.

To rule out nonspecific effects on EPSCs due to pressure application or general inhibitory shunting, we tested local application of glycine (500 µM), another inhibitory transmitter that generates a similarly prominent conductance in SBCs (see Fig. 5 Nerlich et al. 2017), but has no presynaptic effect (see Fig. 2D and E). Glycine did not affect EPSCs (Fig. S6C, D;

$EPSC_{onset}$: control = 1.9 ± 0.3 nA, glycine = 1.9 ± 0.3 nA, $EPSC_{steady}$: control = 1.5 ± 0.3 nA, glycine = 1.6 ± 0.3 nA, n=7, control vs. glycine p = 0.71, two-way RM ANOVA).

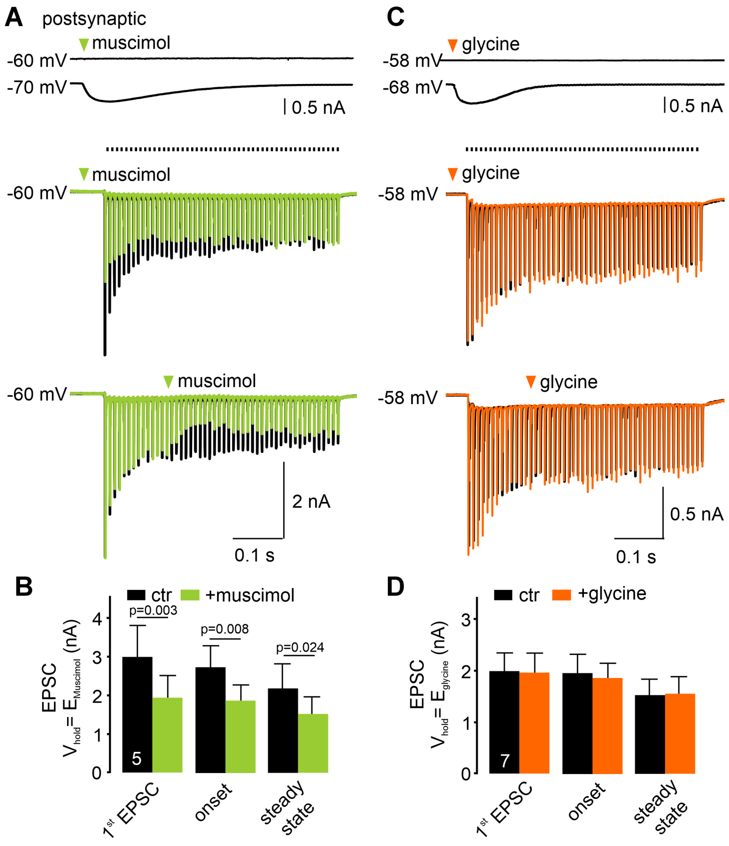


**Fig. S6 Presynaptic GABA$_A$R mediate suppression of glutamatergic transmission**

(A) Postsynaptic currents evoked by muscimol, a selective GABA$_A$R agonist, recorded at $V_{hold}$ = $E_{muscimol}$ (-60 mV) and $V_{hold}$ = -70 mV (top). EPSCs at $E_{muscimol}$ are strongly reduced during muscimol application, consistent with a presynaptic effect.

(B) Mean EPSC amplitudes before and during muscimol application (control vs. muscimol, p = 0.013, n=5, two-way RM ANOVA; Holm–Sidak post hoc).

(C) Postsynaptic currents evoked by glycine puffs are shown at $V_{hold}$ = $E_{glycine}$ (-58 mV) and at -68 mV (top). EPSCs recorded at $E_{glycine}$ are unchanged by glycine, indicating no presynaptic effect of glycine (see also Fig. 2).

(D) EPSCs at $E_{glycine}$ show that GlyR activation in SBCs does not alter EPSC amplitude (control vs. glycine, p = 0.71, n=7, two-way RM ANOVA). Numbers within the bars indicate sample sizes.

Reducing the GABA dose to 100 µM also significantly decreased EPSCs amplitudes and reduced presynaptic action potential and calcium signal amplitudes, confirming a dose-dependent suppression of synaptic transmission (Fig. S7 A-F).

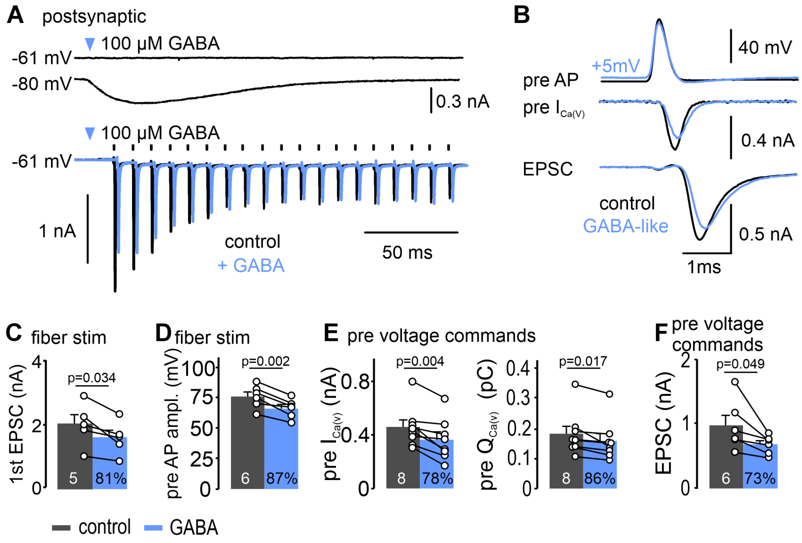


**Fig. S7 Low GABA concentrations suppress glutamatergic transmission by attenuating presynaptic APs and $Ca^{2+}$ influx**

(A) Postsynaptic currents evoked by 100 µM GABA at $V_{hold}$ = $E_{GABA}$ (-61 mV) and -80 mV illustrating the time course of GABA action (top). At $E_{GABA}$, EPSCs amplitudes are reduced during GABA application (bottom).
(B) Presynaptic voltage-clamp recordings using control or GABA-attenuated AP command waveforms (top, see Fig. 5C), with simultaneous presynaptic $I_{Ca(V)}$ (middle) and postsynaptic EPSCs (bottom). GABA-attenuated APs evoke smaller $I_{Ca(V)}$ and EPSCs ($V_{hold}$ pre = -75 mV, post = -70 mV; 1.2 mM external $Ca^{2+}$).
(C-D) EPSC amplitudes evoked by fiber stimulation are reduced by 100 µM GABA (C), due to a smaller presynaptic AP amplitude (D).
(E) GABA-attenuated AP waveforms reduce presynaptic $I_{Ca(V)}$ (left) and total $Ca^{2+}$ charge (right).
(F) EPSCs evoked by GABA-attenuated APs are significantly smaller than controls.
In C-F, numbers in black bars indicate sample sizes; numbers in blue bars give amplitudes normalized to control. *p* values were obtained using paired t-tests.
